# An openly licensed benchmark and per-gene calibration map for missense pathogenicity predictors on activating cancer drivers

**DOI:** 10.64898/2026.07.16.739080

**Authors:** Sung-Gwon Lee

## Abstract

Missense pathogenicity predictors such as AlphaMissense are increasingly used in clinical variant interpretation, yet they are trained on germline labels dominated by loss-of-function (LOF) variants. Using an openly licensed, reproducible benchmark of 768 Cancer Gene Census genes scored with 49 predictors (labels from CIViC, COSMIC, cancerhotspots, ClinVar and gnomAD), we show that 42 of 49 tools (86%) score oncogene, gain-of-function (GOF) variants worse than tumour-suppressor variants. This under-scoring is mechanistically characterized: missed drivers occupy low-conservation, solvent-exposed, non-destabilizing positions (phyloP 2.51 versus 7.89; relative solvent accessibility 0.671 versus 0.185; gene-clustered p = 4.8×10⁻²⁰ and 2.3×10⁻³⁵), and, counter-intuitively, the unsupervised and protein-language models now entering clinical use are the most affected. Per-gene oncogenic thresholds span 0.07–0.99, so a single global cut-off is mis-calibrated for most genes; we provide a per-gene calibration map. A cancer-calibrated stack (OncoCal) modestly improves discrimination over the best single tool (AUROC ≈ 0.93 versus 0.87), rescues drivers such as JAK2 V617F (0.334→0.57), and generalizes to independent deep mutational scanning data. We provide an openly licensed framework to interpret and recalibrate these tools in the somatic setting rather than a replacement predictor.

## Introduction

Computational missense predictors have become central to clinical variant interpretation. Tools such as SIFT, PolyPhen-2, CADD, REVEL, EVE, ESM1b and, most recently, AlphaMissense are widely deployed, and AlphaMissense is being adopted into clinical and research pipelines at scale [1–7]. Almost all these methods are trained or calibrated, directly or indirectly, on germline disease labels (ClinVar, HGMD) with population-frequency proxies for benignity (gnomAD) [3, 5, 8]. Their positive signal therefore reflects a single underlying question — *does this substitution disrupt a conserved or structurally constrained residue?* — which captures loss-of-function (LOF) and destabilizing variants well [9].

Somatic cancer driver biology is different. Activating, gain-of-function (GOF) drivers frequently do not destabilize their protein; they release autoinhibition, create new dimerization interfaces, or allosterically activate signaling, and several of the most important are only mildly disruptive at the residue level [9, 10]. The canonical example is JAK2 V617F, one of the most recurrent point mutations in COSMIC (42,928 samples)[11], which AlphaMissense scores at only 0.334. Prior work has reported this mismatch phenomenologically: germline tools under-perform on GOF or somatic variants [10, 12–14], REVEL reaches strong pathogenic thresholds for fewer GOF than LOF variants [12], driver predictors tend to recognize driver *genes* rather than variant effects [15], and globally calibrated thresholds hide large per-gene heterogeneity [16]. Consistent with this, we have found that in-silico predictors (AlphaMissense, PolyPhen-2) return discordant — and often benign — scores for rare and recurrent JAK/STAT variants, including COSMIC-listed mutations classified benign in ClinVar [17, 18]. Most directly, Tran et al. [19] re-stacked existing scores against a curated oncogenicity knowledgebase and observed that oncogene (GOF) variants are harder to predict than tumor-suppressor variants.

Three gaps remain. First, the *mechanism* of under-scoring — why specific drivers are missed — has not been quantified at scale. Second, the most comprehensive curated oncogenicity resources carry licensing terms that restrict model training and redistribution, limiting reproducibility and reuse. Third, no openly licensed, per-gene calibration of these tools exists for the somatic setting. Here we address all three. Using exclusively open, redistribution-clean labels, we (i) quantify the systematic GOF under-scoring across 49 tools, (ii) characterize it mechanistically through conservation and solvent exposure, (iii) show that newer unsupervised tools are *more* affected than supervised ensembles, (iv) build a cancer-calibrated model with strict anti-circularity controls, (v) provide a per-gene calibration map, and (vi) demonstrate generalization on independent DMS data. We frame this work as a mechanistic characterization and calibration framework, not as a superior predictor.

## Methods

### Gene panel and variant universe

We used the COSMIC Cancer Gene Census (v104, GRCh38) [11, 20] to define 768 cancer genes with oncogene/TSG/fusion role annotations, assigning genes by genomic coordinate interval (CGC GENOME_START–STOP) to recover 48 genes (e.g., BRAF) lacking UniProt cross-references. We enumerated all possible panel missense variants (4,612,188; 747 genes with AlphaMissense scores).

### Labels (open sources only)

Positives were defined as cancer hotspots residues [21], CIViC oncogenic classifications (CC0) [22], or COSMIC recurrence ≥ 10 distinct samples. Negatives were ClinVar benign/likely-benign [23] or gnomAD common variants (AF ≥ 0.001) [24] lacking any positive signal. Variants with simultaneous positive and negative evidence (conflict) or no evidence (unknown) were excluded; unknown variants were never treated as negative. Mechanism (GOF/LOF) used CIViC variant-level annotation where available, otherwise CGC gene-level role as a proxy [25].

### Features

dbNSFP 5.3.1a (GRCh38, BGZF/tabix) [6] provided 49 tool rank scores (0–1; supervised meta-predictors included MutScore [26], VARITY [27], ClinPred [28], BayesDel [29], MetaRNN [30], M-CAP [31], MVP [32], DEOGEN2 [33]), conservation (GERP++ [34], phyloP [35], phastCons [36]; background-selection B-statistic [37]) and gnomAD allele frequencies. Structural features (relative solvent accessibility [RSA], pLDDT) were derived from AlphaFold structures for 734 proteins. RSA was computed with FreeSASA v2.2.1 [38] (Lee–Richards algorithm, 1.4 Å probe, 20 slices, ProtOr radii [39]) on AlphaFold DB (v6) monomer structures [40], as total residue SASA divided by FreeSASA’s per-residue reference maximum (e.g., Ala 108.8, Gly 81.1, Arg 238.2 Å²); residues with RSA < 0.25 were classified as buried. Under this normalization RSA can marginally exceed 1 for fully exposed residues (here, 80 of 3,693 residues; maximum 1.16). Sequence-biochemical features (14; e.g., cysteine gain, charge/hydrophobicity change) were computed from the substitution. All label-defining columns (recurrence, hotspot, CIViC, ClinVar, gnomAD) were excluded from the feature set.

### Join keys

Same-build sources (COSMIC, dbNSFP, AlphaMissense; all GRCh38) were joined on genomic coordinates (chr, pos, ref, alt) because transcript-isoform protein numbering is inconsistent across sources (e.g., BRAF V600E annotated as V640E in genome-screen transcripts). Protein-level keys (GENE:p.<REF><POS><ALT>, one-letter) were used to bridge different-build (CIViC, GRCh37) and residue-based (cancer hotspots) sources.

### Predictor classification

For the supervised-versus-unsupervised analysis, each predictor was classified by whether its training used clinical/pathogenicity labels (supervised; n = 28) or not (unsupervised/self-supervised — conservation, evolutionary, population-genetic or protein-language models; n = 21). Borderline tools (MPC [41], LIST-S2 [42], PrimateAI [43]) were assigned conservatively; because the conservatively assigned high-miss tools (MPC, LIST-S2) fall in the supervised group, the unsupervised-versus-supervised difference is robust to their reassignment. The full classification is provided in S3 Table (type2).

### Model and evaluation

We trained LightGBM classifiers [44] on unique protein-level keys (20,815; 3,922 positive) under StratifiedGroupKFold (5-fold, grouped by gene) with out-of-fold prediction to prevent gene-recognition leakage. Feature ablations were tools-only, tools+mechanism, tools+mechanism+structure. Performance was AUROC and AUPRC, overall and stratified by oncogene/TSG role. Per-gene oncogenic thresholds were derived by Youden’s J; we report both in-sample thresholds and a leave-one-gene-out refinement. Feature attribution used SHAP [45]. We trained OncoCal with default LightGBM hyperparameters across 10 random seeds and report performance as mean ± standard deviation; leave-one-gene-out calibration derived each gene’s threshold from all other genes. Group comparisons of conservation and RSA used two-sided Mann–Whitney U tests; given the few pre-specified comparisons, no multiple-testing correction was applied. Analyses used Python 3.11 with scikit-learn, LightGBM 4.6 and SHAP.

### External validation

15 ProteinGym v1.3 DMS assays [46] were predicted with the trained model; construct numbering was reconciled to AlphaFold canonical sequence by target-sequence alignment (e.g., MET coverage 76 → 1,670 variants). We report Spearman correlation between predictions and experimental DMS fitness for the model, AlphaMissense and REVEL.

### Reproducibility

All code is provided (see Availability of data and materials). COSMIC raw data are not redistributed; only summary statistics and access paths are released.

## Results

### A public, redistribution-clean oncogenicity benchmark

We defined a panel of 768 Cancer Gene Census genes (257 oncogenes, 254 tumor suppressors, 71 with both roles, 140 fusion genes) and enumerated 4,612,188 possible panel missense variants, of which 747 genes carried AlphaMissense scores. Oncogenicity labels were derived exclusively from openly licensed sources: positives from cancer hotspots residues, CIViC oncogenic classifications (CC0), or COSMIC recurrence ≥ 10 samples; negatives from ClinVar benign/likely-benign or gnomAD common variants (allele frequency ≥ 0.001); variants with conflicting or absent evidence were excluded rather than treated as negative. This yielded 20,815 unique protein-level training entries (3,922 positive). COSMIC-derived data are reported only as summary statistics. Each variant was annotated with 49 dbNSFP 5.3.1a tool rank scores, conservation metrics (GERP++, phyloP, phastCons), AlphaFold-derived RSA and pLDDT, and 14 sequence-biochemical features (Methods).

### Predictors systematically under-score activating cancer drivers, and unsupervised tools most of all

Across all 49 tools, 42 (86%) achieved a lower AUROC for oncogene (gain-of-function) variants than for tumor-suppressor (loss-of-function) variants, with a mean gap of 0.030 (sign test p = 3.6×10⁻⁷) (Fig 1A; S1 Table). This is not a property of any single tool: the best performers reached AUROC ≈ 0.87 (MutScore 0.876, VARITY_R 0.874, ClinPred 0.869, AlphaMissense 0.868, BayesDel 0.864), yet the directional bias was near-universal.

**Fig. 1.**
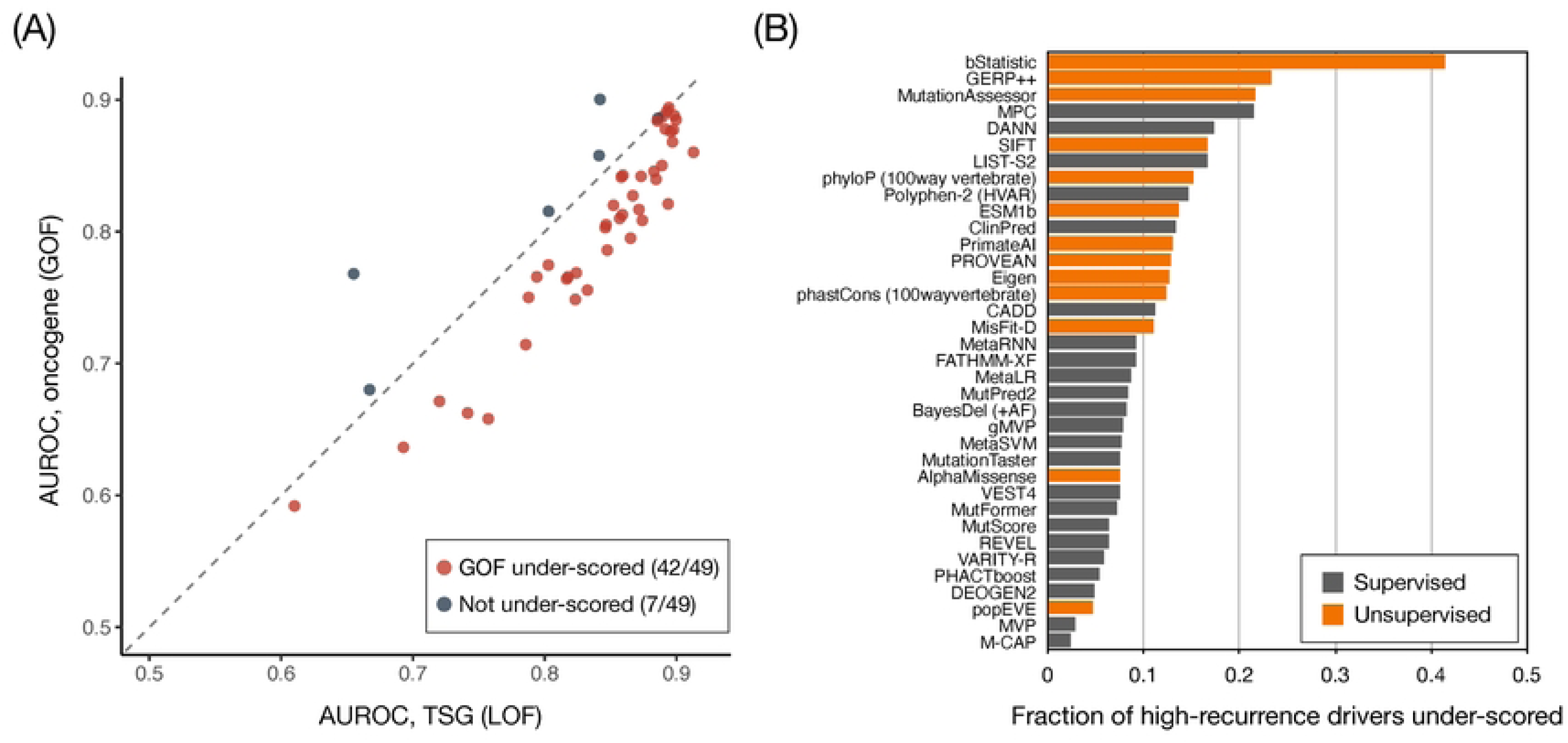
Missense predictors systematically under-score activating cancer drivers — most of all the unsupervised ones. (A) Per-tool AUROC for oncogene (gain-of-function) versus tumor-suppressor (loss-of-function) variants across 49 predictors. Each point is one predictor; red if oncogene AUROC < TSG AUROC (42/49 tools; mean gap 0.030). Dashed line, equality. (B) Fraction of high-recurrence drivers (≥ 50 samples, n = 485) under-scored (rankscore < 0.5) per predictor, colored by class — supervised (clinical-label-trained; n = 28) versus unsupervised/self-supervised (conservation, evolutionary or protein-language models; n = 21). Unsupervised tools under-score 17.8% on average versus 9.1% for supervised tools. One representative predictor per tool family is shown (36 of 49); summary percentages are computed over all 49.

The bias was concentrated by training paradigm. We classified the 49 predictors as supervised (trained on clinical/pathogenicity labels; n = 28) or unsupervised/self-supervised (conservation, evolutionary or protein-language models trained without clinical labels; n = 21). Among high-recurrence drivers (≥ 50 samples, n = 485), unsupervised tools under-scored 17.8% on average versus 9.1% for supervised tools (Fig 1B; S3 Table). The worst offenders were conservation/constraint scores (bStatistic 0.41, GERP++ 0.23 miss rate) and unsupervised language/evolutionary models (ESM1b, EVE); the most robust were supervised meta-predictors (M-CAP 0.024, MVP 0.029, DEOGEN2 0.049). This is counter-intuitive given the strong germline-benchmark performance of unsupervised models and has a practical implication: the very tools now entering clinical use — including AlphaMissense and protein-language models — are among the most prone to missing activating drivers, because their signal is closest to raw evolutionary/structural constraint. A threshold-free comparison confirmed that this is not merely a calibration effect: unsupervised tools had significantly lower oncogene AUROC (0.744 vs 0.835; permutation p = 1×10⁻⁴). The gap was not exclusive to gain-of-function — unsupervised tools were also weaker on tumor-suppressor variants (0.784 vs 0.857) — indicating lower overall discrimination on cancer-driver variants that is most pronounced for, but not specific to, activating drivers.

### AlphaMissense misses a clinically important subset of drivers, characterized by low conservation and surface exposure

We next examined AlphaMissense, the flagship tool now entering clinical pipelines. It failed to reach its pathogenic threshold (raw score < 0.5) for 1,071 of 3,922 unique oncogenic driver positives (27.3%; Fig 2A). Most well-studied drivers were scored correctly, but a clinically important subset was missed despite high recurrence — most strikingly the single most recurrent driver JAK2 V617F (score 0.334; 42,928 samples), together with FGFR3 Y373C (0.175), MPL W515L (0.269) and STAT3 Y640F (0.368); the same tool correctly scored BRAF V600E (0.993; 31,896 samples), KRAS G12D (0.998), EGFR L858R (0.997), IDH1 R132H (0.992) and KIT D816V (0.999). STAT3 Y640F, for instance, lies in the SH2 domain, where activating substitutions drive constitutive signaling [47]. Recurrence alone therefore does not predict whether a driver is scored correctly, pointing to a residue-level determinant. (A full predictor-by-driver comparison across 58 canonical drivers, including tumor-suppressor contrasts, is in Table 1 and S1 Fig; note that S1 Fig shows dbNSFP rank scores — percentile ranks that compress the raw-score scale, so JAK2 V617F appears at rank score 0.59 despite its raw pathogenicity of 0.334.)

**Fig. 2.**
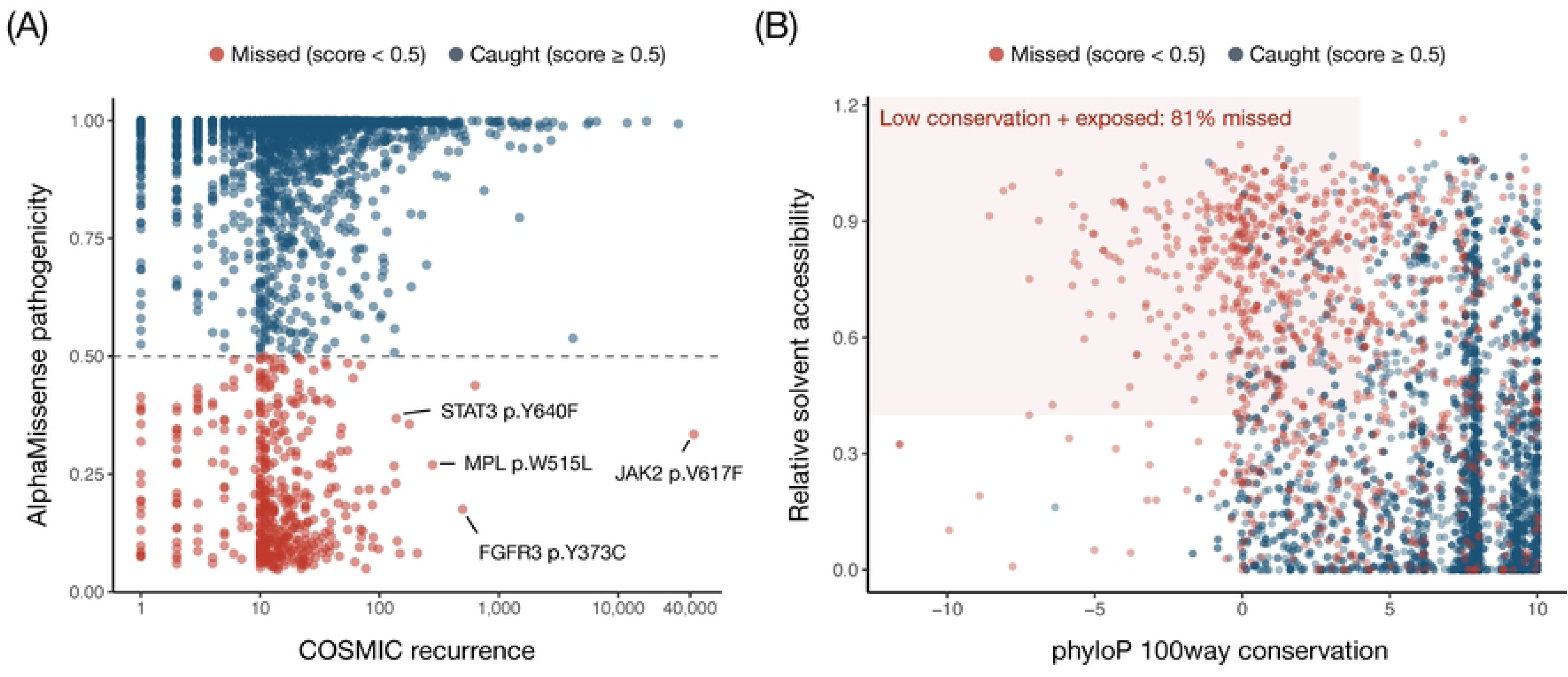
AlphaMissense misses a clinically important subset of drivers, jointly characterized by low conservation and high solvent exposure. (A) AlphaMissense raw pathogenicity versus COSMIC recurrence (log scale) for 3,922 oncogenic driver positives (1,071, 27.3%, AM-missed). Dashed line, pathogenic threshold (0.5); red, below threshold (“missed”, n = 1,071, 27.3%); blue, at or above (“caught”). Labelled, high-recurrence missed drivers (JAK2 V617F, FGFR3 Y373C, MPL W515L, STAT3 Y640F). (B) Conservation (phyloP100way) versus relative solvent accessibility (RSA) for missed (red) and caught (blue) positives. The two axes are largely orthogonal and act jointly: variants in the low-conservation, solvent-exposed corner (phyloP < 4 and RSA > 0.4; shaded) are 81% missed versus 15% elsewhere. Marginal distributions and exact statistics are in S2 Fig.

**Table 1.**
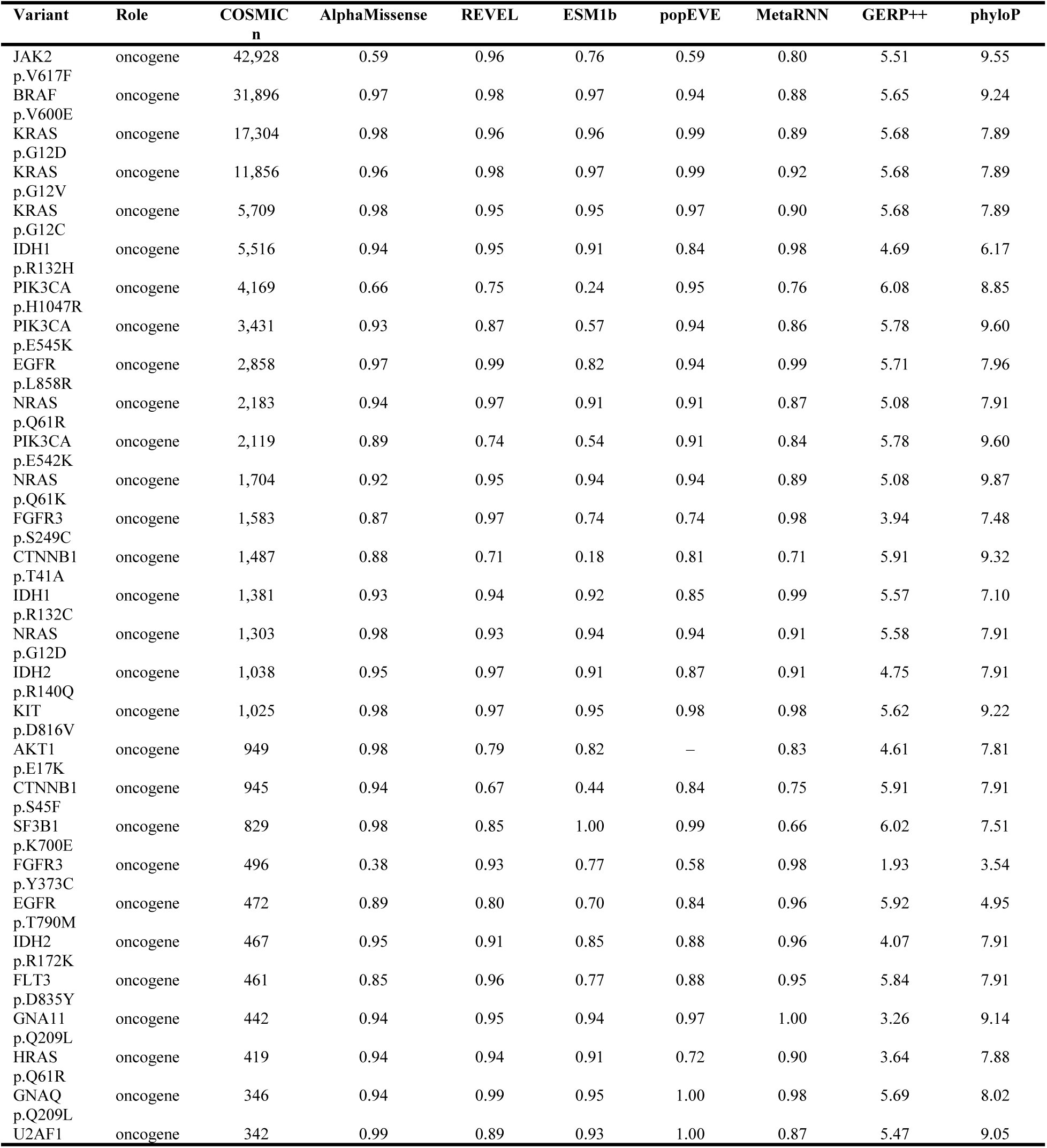
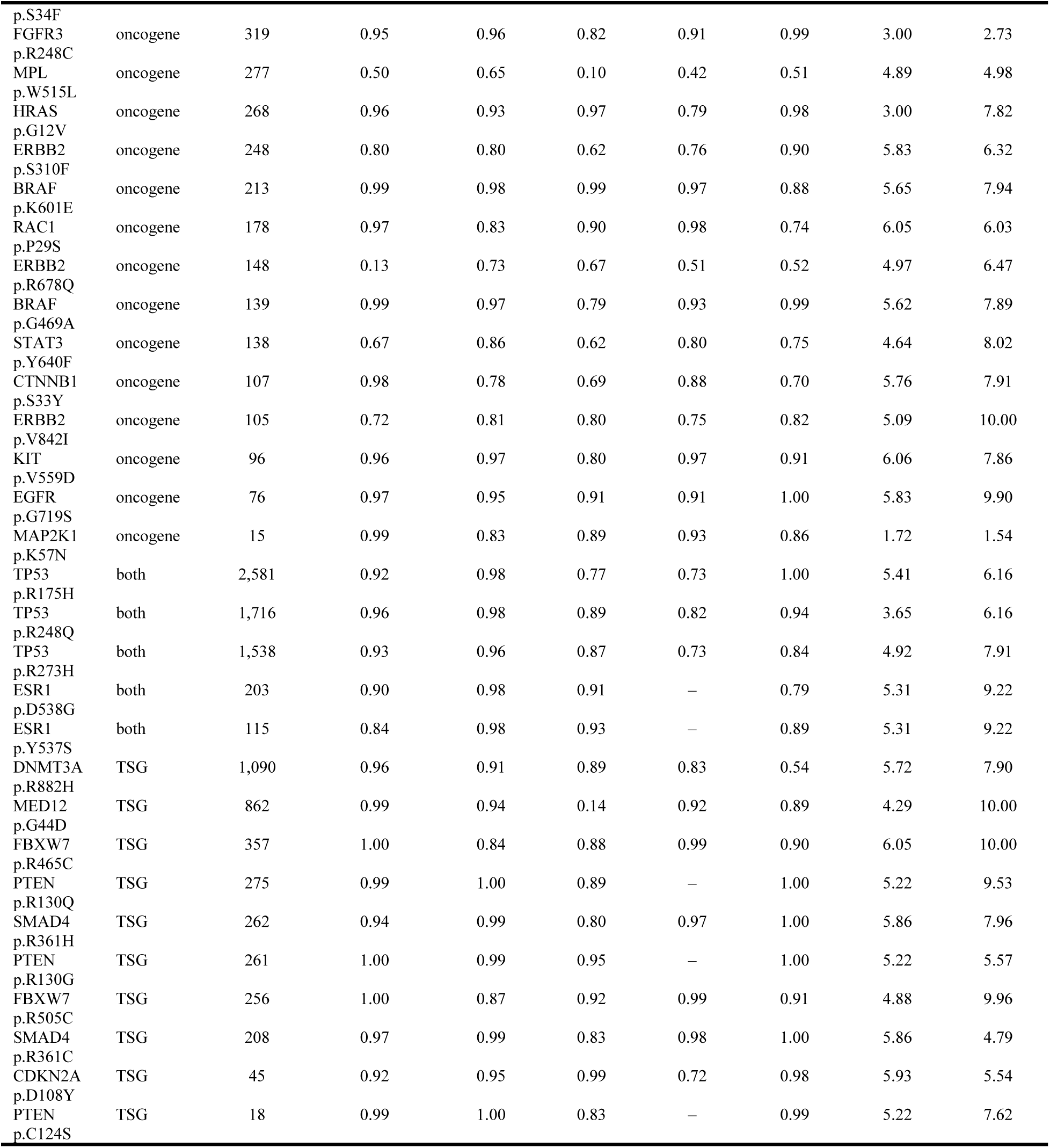
Predictor scores and COSMIC recurrence for canonical cancer driver mutations. For 58 canonical drivers (43 oncogene, 5 dual-role, 10 tumor-suppressor), the protein change, gene role, COSMIC sample count, and scores from representative predictors (AlphaMissense, REVEL, ESM1b, popEVE [56], MetaRNN) and conservation metrics are shown. Activating drivers such as JAK2 V617F receive low scores despite the highest recurrence. Scores are dbNSFP rankscores (percentile ranks).

That determinant is twofold. Comparing the missed and caught oncogenic positives, missed drivers sat at markedly less conserved positions (phyloP100way median 2.51, n = 1,069, vs 7.89, n = 2,851; gene-clustered p = 4.8×10⁻²⁰; GERP++ RS 3.92 vs 5.25) and at more solvent-exposed, non-destabilizing positions (RSA 0.671 vs 0.185; gene-clustered p = 2.3×10⁻³⁵; S2 Table; marginal distributions in S2 Fig). These two axes are largely orthogonal and act jointly: among variants in the low-conservation, solvent-exposed corner (phyloP < 4 and RSA > 0.4) 81% are missed, versus 15% elsewhere (Fig 2B). The tools track residue conservation and structural burial — proxies for LOF damage — rather than oncogenicity itself, a mechanistic instantiation of the gene-recognition critique of driver predictors [15]. Treating each gene as the independent unit to avoid pseudo-replication, a cluster-robust GEE preserved both associations (phyloP OR = 0.47 per SD, 95% CI 0.40–0.55; RSA OR = 2.09, 95% CI 1.86–2.34), and the signal held within single-source (hotspot-only, COSMIC-recurrence-only) and recurrence-matched strata (gene-clustered p ≤ 4×10⁻⁶), excluding an ascertainment artifact.

A residual class escapes even this explanation. JAK2 V617F sits at a highly conserved position (phyloP 9.5) yet is still missed, consistent with a moderate V→F substitution that relieves JH2 pseudokinase autoinhibition [48, 49] — a mechanism invisible to conservation- and stability-based scoring.

### A cancer-calibrated model partially rescues missed drivers, with honest limits

We trained OncoCal — a cancer-calibrated stack of existing predictors — as LightGBM models under gene-group 5-fold cross-validation (out-of-fold predictions, unique protein keys, recurrence excluded as a feature) across 10 random seeds. The model reached AUROC 0.925 ± 0.005 (tools only), 0.930 ± 0.004 (+ mechanism) and 0.924 ± 0.010 (+ mechanism + structure), all far above the best single tool (VARITY_R 0.871; AlphaMissense 0.870) — a margin that greatly exceeds the across-seed standard deviation. The corresponding AUPRC (≈ 0.83) likewise exceeded VARITY_R (0.746) and AlphaMissense (0.759) against a positive-class prevalence of 0.188 (Fig 3; S4 Table). Critically, the three feature sets were statistically indistinguishable (overlapping standard deviations): adding sequence-mechanism and AlphaFold structural features did not reliably improve on the tool-stack-only model, and only structural pLDDT entered the top features among non-tool inputs (top features overall: ClinPred, BayesDel, AlphaMissense, bStatistic, MutScore, structural pLDDT, MetaRNN; S5 Table). ROC and precision-recall curves confirm this ordering (S3 Fig). The model nonetheless rescued ≈ 14% of AlphaMissense-missed oncogenic positives — for example JAK2 V617F (AlphaMissense 0.334 → model 0.57). On bootstrap resampling this advantage was statistically firm: AUROC 0.928 (95% CI 0.923–0.933) versus AlphaMissense 0.870 (0.862–0.878), a gap of 0.058 (95% CI 0.052–0.064) with non-overlapping intervals.

**Fig. 3.**
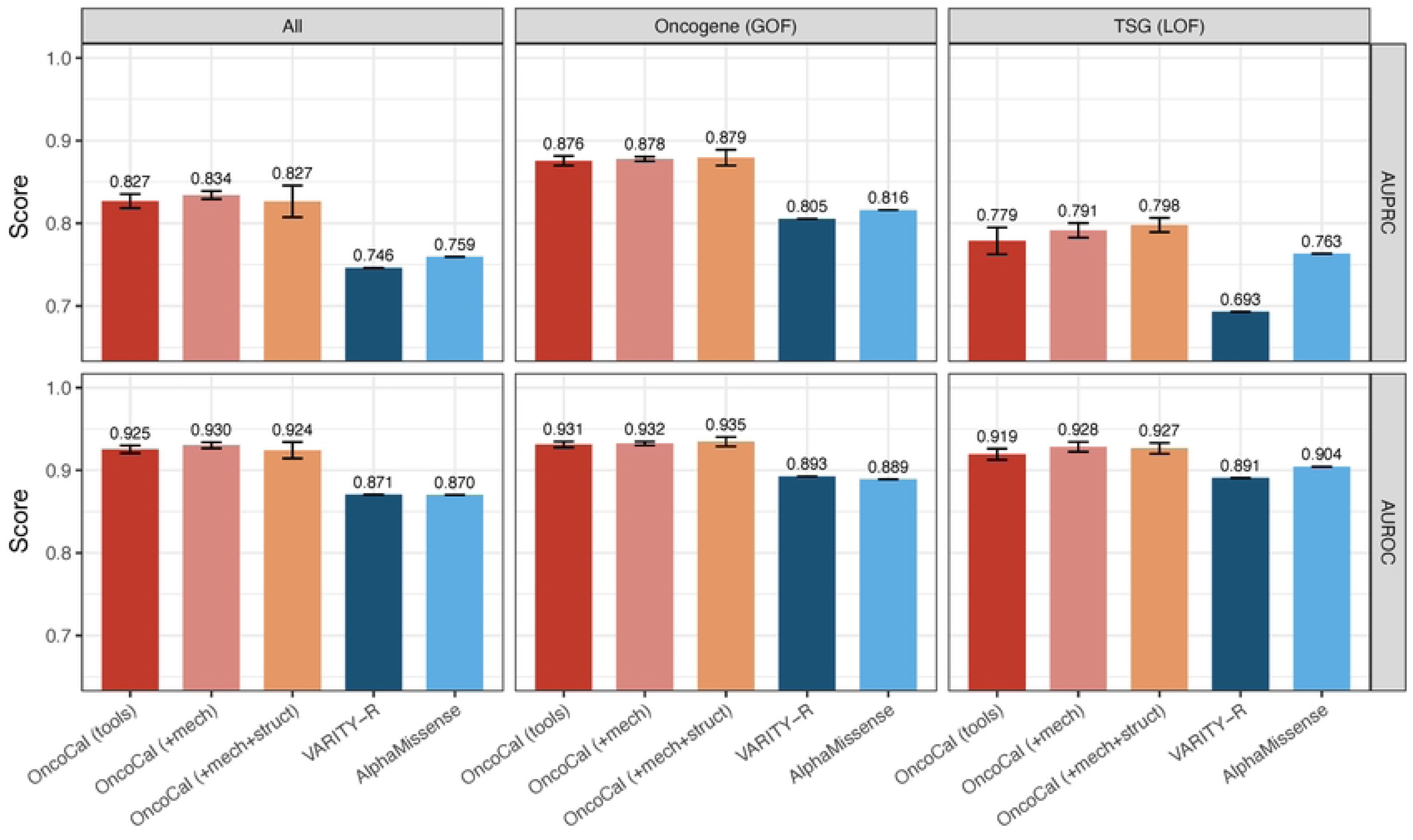
Cancer-calibrated model (OncoCal) versus predictors (gene-group out-of-fold, 10 seeds). AUROC and AUPRC (mean ± s.d. over 10 random seeds; error bars) overall and stratified by oncogene/TSG role, for OncoCal with tools only, + mechanism, and + mechanism + structure features, compared with the best single tool (VARITY-R) and AlphaMissense. OncoCal reaches AUROC ≈ 0.93 / AUPRC ≈ 0.83 versus ≈ 0.87 / ≈ 0.75 for the best single tool — a margin far exceeding the seed s.d. — whereas the three feature sets overlap within s.d., showing that mechanism/structure features add no reliable gain. Dotted line in AUPRC panels marks the positive prevalence (0.188), the random-classifier baseline.

We state the interpretation plainly. The model is fundamentally a stack of existing tools; its value is stack-level integration and calibrated rescue of specific drivers, not a new mechanistic signal or a categorical accuracy gain. Mechanism and structural features add little because their information is already absorbed by the constituent predictors. We therefore do not claim that mechanism/structure features improve prediction. Because its constituent scores were themselves trained or calibrated on germline benign labels that overlap our negative set, the absolute AUROC margin over the best single tool is likely optimistic and should be read as stack-level integration rather than new generalization.

### Per-gene calibration reveals clinically actionable heterogeneity

For 118 genes with sufficient labels, the AlphaMissense score threshold separating oncogenic from non-oncogenic variants ranged from 0.071 to 0.999 (median 0.277, IQR 0.185–0.582) — far wider than the single global Youden threshold of 0.469 (Fig 4; S6 Table). For example, LRP1B (TSG) reached its optimal cut at AlphaMissense ≥ 0.09, whereas SMAD4 required ≥ 0.99 and SMARCB1 ≥ 0.999. Applying per-gene rather than global thresholds improved balanced accuracy, under within-gene leave-one-out, from 0.813 to 0.829 (+0.016) for AlphaMissense and from 0.843 to 0.874 (+0.032) for the model (in-sample gains, +0.036 and +0.047, are optimistic). This is the somatic-oncogenicity analogue of the germline per-gene calibration heterogeneity reported by Tejura et al. [16], and yields a directly usable clinical map (“for this gene, a score ≥ T indicates oncogenicity”). The leave-one-out procedure that removes in-sample optimism is detailed in Methods and Limitations.

**Fig. 4.**
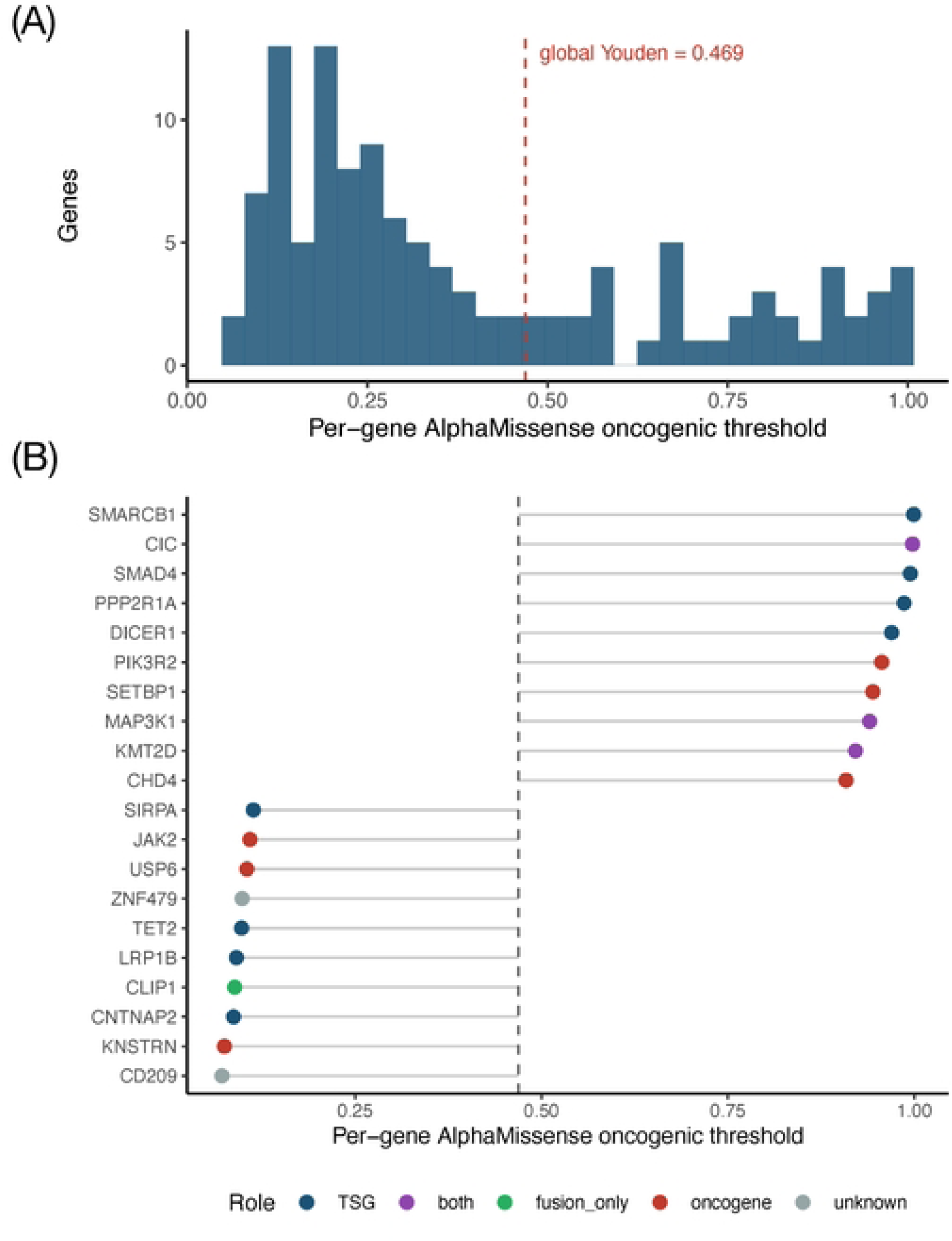
Per-gene oncogenic thresholds reveal clinically actionable heterogeneity. (A) Distribution of per-gene AlphaMissense oncogenic thresholds (n = 118 genes), spanning 0.07–0.99 (median 0.277) versus the single global Youden threshold (0.469, dashed line). (B) Forest plot of genes requiring the lowest and highest thresholds, colored by gene role.

### Independent validation on deep mutational scanning data

To test generalization beyond our label scheme, we predicted independent DMS measurements from ProteinGym v1.3 (15 assays) using the full model and correlated predictions with experimental fitness (Fig 5; S8 Table). The model achieved |Spearman| 0.492, comparable to AlphaMissense (0.502) and better than REVEL (0.426); all correlations had the correct sign, and the model outperformed AlphaMissense on 6 of 15 assays (e.g., PPARG model −0.737 vs −0.616; BRCA1 −0.590; PPM1D −0.626; MET −0.544 vs −0.580; AlphaMissense led on SRC across four assays). Importantly, GOF superiority is *not* visible in global DMS correlation, because global DMS rank correlation is dominated by the LOF/neutral majority even in nominally activating assays. GOF rescue is a property of rare, recurrent drivers and is demonstrated through the canonical-driver and recurrence analyzes (above), not through global DMS correlation; we state this explicitly as a limitation.

**Fig. 5.**
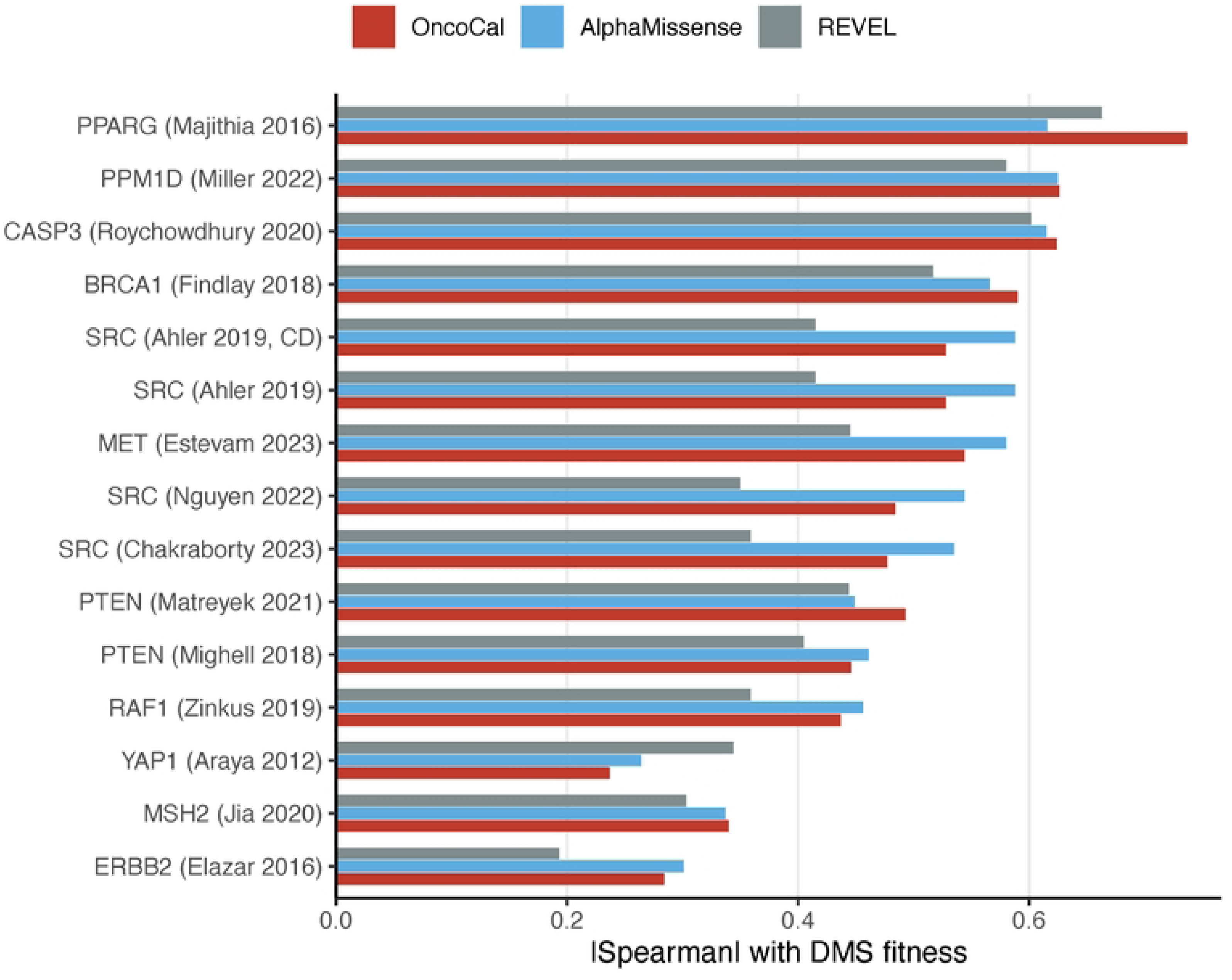
Generalization to independent deep mutational scanning data. Absolute Spearman correlation between predictions and experimental DMS fitness across 15 ProteinGym v1.3 assays for OncoCal, AlphaMissense and REVEL. OncoCal (|Spearman| 0.492) is comparable to AlphaMissense (0.502) and exceeds REVEL (0.426); all correlations have the correct sign.

## Discussion

Germline missense predictors systematically under-score activating cancer drivers, and we show this is mechanistically characterized: missed drivers sit at low-conservation and/or solvent-exposed, non-destabilizing positions, because the tools effectively measure residue constraint and structural damage — hallmarks of LOF — rather than oncogenic activation. The phenomenon is near-universal (42/49 tools) and, notably, is *worse* for unsupervised/self-supervised predictors (conservation, evolutionary and protein-language models, including AlphaMissense; 17.8% vs 9.1%) than for supervised tools. As fields migrate to AlphaMissense and related models, this is an actionable caution: a low score in a known oncogene, particularly at a poorly conserved or surface residue, should not be read as evidence against oncogenicity. More broadly, accurate interpretation of activating variants is clinically consequential: rare and recurrent JAK/STAT variants can exert substantial functional and transcriptomic effects even when germline-trained predictors score them benign [17, 18, 50]. Concurrent work using a knowledge-based distance metric similarly reports that variant effect predictors underperform on gain-of-function variants, which often exert milder, less destabilizing effects than conservation-based scoring assumes [51].

Our contribution is deliberately framed as characterization and calibration rather than a new predictor. Cancer-specific driver predictors such as boostDM [52] and CHASMplus [53] exist, but the tools deployed in clinical variant interpretation are germline-trained (AlphaMissense, REVEL); our aim is to characterize and recalibrate those rather than replace them. The cancer-calibrated model rescues specific drivers (JAK2 V617F, ≈14% of AlphaMissense-missed positives) but offers only incremental discrimination because it is a tool stack, and structural features (RSA and pLDDT) add little beyond what the tools already encode. The more durable outputs are (i) the mechanistic explanation, (ii) the per-gene calibration map, which converts a single mis-calibrated global threshold into gene-specific, clinically usable cut-offs, and (iii) an openly licensed benchmark that any group can reproduce and extend. A recent interpretable framework, MutaPheno, likewise reports that germline pathogenicity tools lose accuracy on cancer drivers and builds a mechanism-grounded classifier from structural and biophysical features [54]; our work is complementary, contributing the unsupervised-versus-supervised dissociation and an openly licensed per-gene somatic calibration map rather than a new classifier.

This study differs from the closest prior work, Tran et al. (2025) [19], in five concrete ways. Tran et al. re-stacked existing scores against a curated oncogenicity knowledgebase and *observed* the oncogene gap; we instead (1) *characterize* the gap mechanistically via conservation and solvent exposure, (2) decompose it into an unsupervised-versus-supervised dissociation, (3) use exclusively openly licensed labels, so the resource is reproducible and license-clean, (4) provide a per-gene calibration map, and (5) impose explicit anti-circularity controls (gene-group cross-validation, recurrence excluded as a feature). In short, our work is about *characterizing and calibrating* the failure mode rather than re-stacking around it.

### Limitations

Several limitations qualify these findings. OncoCal is fundamentally a stack of existing predictors: its discrimination gain over the best single tool is modest, and adding mechanism or structural features did not reliably improve on the tool-stack baseline (the difference between feature sets fell within cross-validation seed-level variation), so we do not claim a categorically better predictor or that mechanism/structure features improve prediction. Variant-level gain-of-function/loss-of-function mechanism prediction was not achievable: available labels were too few (49 CIViC mechanism annotations; 28 gain-of-function and 21 loss-of-function across 17 genes), candidate features did not separate the two classes (all comparisons p > 0.13; S7 Table), and leave-one-gene-out classification (AUROC 0.679) fell far below a gene-role baseline (0.957), indicating that apparent mechanism prediction collapses to gene identity; we therefore provide only gene-level mechanism annotation; dedicated mode-of-action predictors such as PreMode instead pursue variant-level gain/loss classification with larger, structure-aware graph models [55]. Label circularity is mitigated but not eliminated, as positives are recurrence- and hotspot-weighted; although recurrence was excluded as a feature and gene-group splits were used, residual circularity remains. Negatives are germline benign or common variants (ClinVar/gnomAD), which differ in ascertainment from somatic positives. Finally, the deep mutational scanning validation is loss-of-function-weighted: global DMS correlation is dominated by loss-of-function and neutral variants and does not directly demonstrate a gain-of-function advantage, activating-readout assays are scarce, and 26 very large proteins lacked AlphaFold structures; per-gene thresholds are in-sample and require leave-one-gene-out confirmation.

## Conclusions

Germline missense predictors systematically under-score activating cancer drivers — a bias that is characterized by low conservation and high solvent exposure, is worst for the unsupervised tools now entering clinical use and is only partially correctable by stacking. Rather than a replacement predictor, we provide an openly licensed, reproducible benchmark and a per-gene calibration map that allow existing tools to be interpreted and recalibrated in the somatic setting. In practice, a low score from a germline-trained predictor — especially at a poorly conserved or surface residue of a known oncogene — should not be read as evidence against oncogenicity.

## List of abbreviations

AF: allele frequency
AUPRC: area under the precision–recall curve
AUROC: area under the receiver operating characteristic curve
CC0: Creative Commons Zero (public-domain dedication)
CGC: Cancer Gene Census
DMS: deep mutational scanning
GOF: gain-of-function
LOF: loss-of-function
pLDDT: predicted local distance difference test
RSA: relative solvent accessibility
s.d.: standard deviation
SHAP: SHapley Additive exPlanations
TSG: tumor-suppressor gene.

## Declarations

### Ethics approval and consent to participate

Not applicable; this study used only publicly available, de-identified data.

### Availability of data and materials

All analysis code is released on GitHub (https://github.com/tjdrnjsqpf/oncocal; archived at Zenodo, DOI 10.5281/zenodo.20741850). Openly licensed labels and result tables (derived from CIViC [CC0], ClinVar, cancerhotspots, gnomAD) are released with the paper. COSMIC source data are not redistributed in accordance with its academic license; only summary statistics and access instructions are provided. dbNSFP 5.3.1a, AlphaFold and ProteinGym are available from their original providers.

### Authors’ contributions

S.-G.L. is the sole author: conceived and designed the study, performed all analyses, and wrote the manuscript. The author read and approved the final manuscript.

## Acknowledgements

The author thanks Professor Chungoo Park, Dr. Seongmin Kim (Chonnam National University) and Dr. Lothar Hennighausen (National Institutes of Health) for helpful discussions.

## References

1. Adzhubei IA, Schmidt S, Peshkin L, Ramensky VE, Gerasimova A, Bork P, et al. A method and server for predicting damaging missense mutations. Nat Methods. 2010;7(4):248–9. doi: 10.1038/nmeth0410-248. PubMed PMID: 20354512; PubMed Central PMCID: PMCPMC2855889.

2. Brandes N, Goldman G, Wang CH, Ye CJ, Ntranos V. Genome-wide prediction of disease variant effects with a deep protein language model. Nat Genet. 2023;55(9):1512–22. Epub 20230810. doi: 10.1038/s41588-023-01465-0. PubMed PMID: 37563329; PubMed Central PMCID: PMCPMC10484790.

3. Cheng J, Novati G, Pan J, Bycroft C, Zemgulyte A, Applebaum T, et al. Accurate proteome-wide missense variant effect prediction with AlphaMissense. Science. 2023;381(6664):eadg7492. Epub 20230922. doi: 10.1126/science.adg7492. PubMed PMID: 37733863.

4. Frazer J, Notin P, Dias M, Gomez A, Min JK, Brock K, et al. Disease variant prediction with deep generative models of evolutionary data. Nature. 2021;599(7883):91–5. Epub 20211027. doi: 10.1038/s41586-021-04043-8. PubMed PMID: 34707284.

5. Ioannidis NM, Rothstein JH, Pejaver V, Middha S, McDonnell SK, Baheti S, et al. REVEL: An Ensemble Method for Predicting the Pathogenicity of Rare Missense Variants. Am J Hum Genet. 2016;99(4):877–85. Epub 20160922. doi: 10.1016/j.ajhg.2016.08.016. PubMed PMID: 27666373; PubMed Central PMCID: PMCPMC5065685.

6. Liu X, Li C, Mou C, Dong Y, Tu Y. dbNSFP v4: a comprehensive database of transcript-specific functional predictions and annotations for human nonsynonymous and splice-site SNVs. Genome Med. 2020;12(1):103. Epub 20201202. doi: 10.1186/s13073-020-00803-9. PubMed PMID: 33261662; PubMed Central PMCID: PMCPMC7709417.

7. Rentzsch P, Witten D, Cooper GM, Shendure J, Kircher M. CADD: predicting the deleteriousness of variants throughout the human genome. Nucleic Acids Res. 2019;47(D1):D886–D94. doi: 10.1093/nar/gky1016. PubMed PMID: 30371827; PubMed Central PMCID: PMCPMC6323892.

8. Pejaver V, Byrne AB, Feng BJ, Pagel KA, Mooney SD, Karchin R, et al. Calibration of computational tools for missense variant pathogenicity classification and ClinGen recommendations for PP3/BP4 criteria. Am J Hum Genet. 2022;109(12):2163–77. Epub 20221121. doi: 10.1016/j.ajhg.2022.10.013. PubMed PMID: 36413997; PubMed Central PMCID: PMCPMC9748256.

9. Gerasimavicius L, Livesey BJ, Marsh JA. Loss-of-function, gain-of-function and dominant-negative mutations have profoundly different effects on protein structure. Nat Commun. 2022;13(1):3895. Epub 20220706. doi: 10.1038/s41467-022-31686-6. PubMed PMID: 35794153; PubMed Central PMCID: PMCPMC9259657.

10. Ostroverkhova D, Przytycka TM, Panchenko AR. Cancer driver mutations: predictions and reality. Trends Mol Med. 2023;29(7):554–66. Epub 20230417. doi: 10.1016/j.molmed.2023.03.007. PubMed PMID: 37076339; PubMed Central PMCID: PMCPMC12232956.

11. Tate JG, Bamford S, Jubb HC, Sondka Z, Beare DM, Bindal N, et al. COSMIC: the Catalogue Of Somatic Mutations In Cancer. Nucleic Acids Res. 2019;47(D1):D941–D7. doi: 10.1093/nar/gky1015. PubMed PMID: 30371878; PubMed Central PMCID: PMCPMC6323903.

12. Hopkins JJ, Wakeling MN, Johnson MB, Flanagan SE, Laver TW. REVEL Is Better at Predicting Pathogenicity of Loss-of-Function than Gain-of-Function Variants. Hum Mutat. 2023;2023:8857940. Epub 20231204. doi: 10.1155/2023/8857940. PubMed PMID: 40225155; PubMed Central PMCID: PMCPMC11918948.

13. Ostroverkhova D, Sheng Y, Panchenko A. Are Next-Generation Pathogenicity Predictors Applicable to Cancer? J Mol Biol. 2024;436(16):168644. Epub 20240605. doi: 10.1016/j.jmb.2024.168644. PubMed PMID: 38848867.

14. Stein D, Kars ME, Wu Y, Bayrak CS, Stenson PD, Cooper DN, et al. Genome-wide prediction of pathogenic gain- and loss-of-function variants from ensemble learning of a diverse feature set. Genome Med. 2023;15(1):103. Epub 20231130. doi: 10.1186/s13073-023-01261-9. PubMed PMID: 38037155; PubMed Central PMCID: PMCPMC10688473.

15. Raimondi D, Passemiers A, Fariselli P, Moreau Y. Current cancer driver variant predictors learn to recognize driver genes instead of functional variants. BMC Biol. 2021;19(1):3. Epub 20210113. doi: 10.1186/s12915-020-00930-0. PubMed PMID: 33441128; PubMed Central PMCID: PMCPMC7807764.

16. Tejura M, Fayer S, McEwen AE, Flynn J, Starita LM, Fowler DM. Calibration of variant effect predictors on genome-wide data masks heterogeneous performance across genes. Am J Hum Genet. 2024;111(9):2031–43. Epub 20240821. doi: 10.1016/j.ajhg.2024.07.018. PubMed PMID: 39173626; PubMed Central PMCID: PMCPMC11393694.

17. Hennighausen L, Haikarainen T, Lee SG, Caf Y, Furth PA, Silvennoinen O, et al. Investigation of the transcriptional impact of rare germline JAK/STAT variants found in a Tyrolean alpine community. BMC Genomics. 2025;27(1):8. Epub 20251201. doi: 10.1186/s12864-025-12307-0. PubMed PMID: 41327031; PubMed Central PMCID: PMCPMC12771743.

18. Lee HK, Cho G, Huh JW, Furth PA, Kim J, Hennighausen L. Rare germline JAK/STAT variants in a Korean cohort amplify innate immune responses to vaccination. iScience. 2025;28(12):114110. Epub 20251119. doi: 10.1016/j.isci.2025.114110. PubMed PMID: 41467183; PubMed Central PMCID: PMCPMC12744258.

19. Tran TN, Fong C, Pichotta K, Luthra A, Shen R, Chen Y, et al. AI cancer driver mutation predictions are valid in real-world data. Nat Commun. 2025;16(1):8509. Epub 20250926. doi: 10.1038/s41467-025-63461-8. PubMed PMID: 41006257; PubMed Central PMCID: PMCPMC12474978.

20. Sondka Z, Bamford S, Cole CG, Ward SA, Dunham I, Forbes SA. The COSMIC Cancer Gene Census: describing genetic dysfunction across all human cancers. Nat Rev Cancer. 2018;18(11):696–705. doi: 10.1038/s41568-018-0060-1. PubMed PMID: 30293088; PubMed Central PMCID: PMCPMC6450507.

21. Chang MT, Bhattarai TS, Schram AM, Bielski CM, Donoghue MTA, Jonsson P, et al. Accelerating Discovery of Functional Mutant Alleles in Cancer. Cancer Discov. 2018;8(2):174–83. Epub 20171215. doi: 10.1158/2159-8290.CD-17-0321. PubMed PMID: 29247016; PubMed Central PMCID: PMCPMC5809279.

22. Griffith M, Spies NC, Krysiak K, McMichael JF, Coffman AC, Danos AM, et al. CIViC is a community knowledgebase for expert crowdsourcing the clinical interpretation of variants in cancer. Nat Genet. 2017;49(2):170–4. doi: 10.1038/ng.3774. PubMed PMID: 28138153; PubMed Central PMCID: PMCPMC5367263.

23. Landrum MJ, Lee JM, Benson M, Brown GR, Chao C, Chitipiralla S, et al. ClinVar: improving access to variant interpretations and supporting evidence. Nucleic Acids Res. 2018;46(D1):D1062–D7. doi: 10.1093/nar/gkx1153. PubMed PMID: 29165669; PubMed Central PMCID: PMCPMC5753237.

24. Karczewski KJ, Francioli LC, Tiao G, Cummings BB, Alfoldi J, Wang Q, et al. The mutational constraint spectrum quantified from variation in 141,456 humans. Nature. 2020;581(7809):434–43. Epub 20200527. doi: 10.1038/s41586-020-2308-7. PubMed PMID: 32461654; PubMed Central PMCID: PMCPMC7334197.

25. Vogelstein B, Papadopoulos N, Velculescu VE, Zhou S, Diaz LA, Jr., Kinzler KW. Cancer genome landscapes. Science. 2013;339(6127):1546–58. doi: 10.1126/science.1235122. PubMed PMID: 23539594; PubMed Central PMCID: PMCPMC3749880.

26. Quinodoz M, Peter VG, Cisarova K, Royer-Bertrand B, Stenson PD, Cooper DN, et al. Analysis of missense variants in the human genome reveals widespread gene-specific clustering and improves prediction of pathogenicity. Am J Hum Genet. 2022;109(3):457–70. Epub 20220203. doi: 10.1016/j.ajhg.2022.01.006. PubMed PMID: 35120630; PubMed Central PMCID: PMCPMC8948164.

27. Wu Y, Li R, Sun S, Weile J, Roth FP. Improved pathogenicity prediction for rare human missense variants. Am J Hum Genet. 2021;108(10):1891–906. Epub 20210921. doi: 10.1016/j.ajhg.2021.08.012. PubMed PMID: 34551312; PubMed Central PMCID: PMCPMC8546039.

28. Alirezaie N, Kernohan KD, Hartley T, Majewski J, Hocking TD. ClinPred: Prediction Tool to Identify Disease-Relevant Nonsynonymous Single-Nucleotide Variants. Am J Hum Genet. 2018;103(4):474–83. Epub 20180913. doi: 10.1016/j.ajhg.2018.08.005. PubMed PMID: 30220433; PubMed Central PMCID: PMCPMC6174354.

29. Feng BJ. PERCH: A Unified Framework for Disease Gene Prioritization. Hum Mutat. 2017;38(3):243–51. Epub 20170128. doi: 10.1002/humu.23158. PubMed PMID: 27995669; PubMed Central PMCID: PMCPMC5299048.

30. Li C, Zhi D, Wang K, Liu X. MetaRNN: differentiating rare pathogenic and rare benign missense SNVs and InDels using deep learning. Genome Med. 2022;14(1):115. Epub 20221008. doi: 10.1186/s13073-022-01120-z. PubMed PMID: 36209109; PubMed Central PMCID: PMCPMC9548151.

31. Jagadeesh KA, Wenger AM, Berger MJ, Guturu H, Stenson PD, Cooper DN, et al. M-CAP eliminates a majority of variants of uncertain significance in clinical exomes at high sensitivity. Nat Genet. 2016;48(12):1581–6. Epub 20161024. doi: 10.1038/ng.3703. PubMed PMID: 27776117.

32. Qi H, Zhang H, Zhao Y, Chen C, Long JJ, Chung WK, et al. MVP predicts the pathogenicity of missense variants by deep learning. Nat Commun. 2021;12(1):510. Epub 20210121. doi: 10.1038/s41467-020-20847-0. PubMed PMID: 33479230; PubMed Central PMCID: PMCPMC7820281.

33. Raimondi D, Tanyalcin I, Ferte J, Gazzo A, Orlando G, Lenaerts T, et al. DEOGEN2: prediction and interactive visualization of single amino acid variant deleteriousness in human proteins. Nucleic Acids Res. 2017;45(W1):W201–W6. doi: 10.1093/nar/gkx390. PubMed PMID: 28498993; PubMed Central PMCID: PMCPMC5570203.

34. Davydov EV, Goode DL, Sirota M, Cooper GM, Sidow A, Batzoglou S. Identifying a high fraction of the human genome to be under selective constraint using GERP++. PLoS Comput Biol. 2010;6(12):e1001025. Epub 20101202. doi: 10.1371/journal.pcbi.1001025. PubMed PMID: 21152010; PubMed Central PMCID: PMCPMC2996323.

35. Pollard KS, Hubisz MJ, Rosenbloom KR, Siepel A. Detection of nonneutral substitution rates on mammalian phylogenies. Genome Res. 2010;20(1):110–21. Epub 20091026. doi: 10.1101/gr.097857.109. PubMed PMID: 19858363; PubMed Central PMCID: PMCPMC2798823.

36. Siepel A, Bejerano G, Pedersen JS, Hinrichs AS, Hou M, Rosenbloom K, et al. Evolutionarily conserved elements in vertebrate, insect, worm, and yeast genomes. Genome Res. 2005;15(8):1034–50. Epub 20050715. doi: 10.1101/gr.3715005. PubMed PMID: 16024819; PubMed Central PMCID: PMCPMC1182216.

37. McVicker G, Gordon D, Davis C, Green P. Widespread genomic signatures of natural selection in hominid evolution. PLoS Genet. 2009;5(5):e1000471. Epub 20090508. doi: 10.1371/journal.pgen.1000471. PubMed PMID: 19424416; PubMed Central PMCID: PMCPMC2669884.

38. Mitternacht S. FreeSASA: An open source C library for solvent accessible surface area calculations. F1000Res. 2016;5:189. Epub 20160218. doi: 10.12688/f1000research.7931.1. PubMed PMID: 26973785; PubMed Central PMCID: PMCPMC4776673.

39. Tsai J, Taylor R, Chothia C, Gerstein M. The packing density in proteins: standard radii and volumes. J Mol Biol. 1999;290(1):253–66. doi: 10.1006/jmbi.1999.2829. PubMed PMID: 10388571.

40. Jumper J, Evans R, Pritzel A, Green T, Figurnov M, Ronneberger O, et al. Highly accurate protein structure prediction with AlphaFold. Nature. 2021;596(7873):583–9. Epub 20210715. doi: 10.1038/s41586-021-03819-2. PubMed PMID: 34265844; PubMed Central PMCID: PMCPMC8371605.

41. Samocha KE, Kosmicki JA, Karczewski KJ, O’Donnell-Luria AH, Pierce-Hoffman E, MacArthur DG, et al. Regional missense constraint improves variant deleteriousness prediction. bioRxiv. 2017:148353. doi: 10.1101/148353.

42. Malhis N, Jacobson M, Jones SJM, Gsponer J. LIST-S2: taxonomy based sorting of deleterious missense mutations across species. Nucleic Acids Res. 2020;48(W1):W154–W61. doi: 10.1093/nar/gkaa288. PubMed PMID: 32352516; PubMed Central PMCID: PMCPMC7319545.

43. Sundaram L, Gao H, Padigepati SR, McRae JF, Li Y, Kosmicki JA, et al. Predicting the clinical impact of human mutation with deep neural networks. Nat Genet. 2018;50(8):1161–70. Epub 20180723. doi: 10.1038/s41588-018-0167-z. PubMed PMID: 30038395; PubMed Central PMCID: PMCPMC6237276.

44. Ke G, Meng Q, Finley T, Wang T, Chen W, Ma W, Ye Q, Liu TY. LightGBM: A Highly Efficient Gradient Boosting Decision Tree. Advances in Neural Information Processing Systems (NeurIPS) 302017. p. 3146–54.

45. Lundberg SML, Su-In. A Unified Approach to Interpreting Model Predictions. Advances in Neural Information Processing Systems (NeurIPS) 30: Curran Associates; 2017. p. 4765–74.

46. Notin P, Kollasch A, Ritter D, van Niekerk L, Paul S, Spinner H, Rollins N, Shaw A, Orenbuch R, Weitzman R, Frazer J, Dias M, Franceschi D, Gal Y, Marks DS. ProteinGym: Large-Scale Benchmarks for Protein Fitness Prediction and Design. Advances in Neural Information Processing Systems (NeurIPS) 36, Datasets and Benchmarks Track 2023.

47. Lee HK, Cho G, Chen J, Schultz AB, Lee SG, Liu C, et al. STAT3 SH2 Domain Aspartic Acid 661 Mutations Activate Immune Gene Programs. J Cell Mol Med. 2026;30(1):e71015. doi: 10.1111/jcmm.71015. PubMed PMID: 41527523; PubMed Central PMCID: PMCPMC12796846.

48. Bandaranayake RM, Ungureanu D, Shan Y, Shaw DE, Silvennoinen O, Hubbard SR. Crystal structures of the JAK2 pseudokinase domain and the pathogenic mutant V617F. Nat Struct Mol Biol. 2012;19(8):754–9. Epub 20120722. doi: 10.1038/nsmb.2348. PubMed PMID: 22820988; PubMed Central PMCID: PMCPMC3414675.

49. James C, Ugo V, Le Couedic JP, Staerk J, Delhommeau F, Lacout C, et al. A unique clonal JAK2 mutation leading to constitutive signalling causes polycythaemia vera. Nature. 2005;434(7037):1144–8. doi: 10.1038/nature03546. PubMed PMID: 15793561.

50. Hennighausen L, Cho G, Lee SG, Caf Y, Kim J, Knabl L, et al. Impact of rare JAK/STAT germline mutations on vaccination-induced innate immune responses in a Tyrolian population. Int J Biol Sci. 2026;22(2):519–28. Epub 20260101. doi: 10.7150/ijbs.124098. PubMed PMID: 41522351; PubMed Central PMCID: PMCPMC12780842.

51. Gerasimavicius L, Marsh JA. A knowledge-based distance metric highlights underperformance of variant effect predictors on gain-of-function missense variants. bioRxiv. 2025:2025.07.23.666325. doi: 10.1101/2025.07.23.666325.

52. Muinos F, Martinez-Jimenez F, Pich O, Gonzalez-Perez A, Lopez-Bigas N. In silico saturation mutagenesis of cancer genes. Nature. 2021;596(7872):428–32. Epub 20210728. doi: 10.1038/s41586-021-03771-1. PubMed PMID: 34321661.

53. Tokheim C, Karchin R. CHASMplus Reveals the Scope of Somatic Missense Mutations Driving Human Cancers. Cell Syst. 2019;9(1):9–23 e8. Epub 20190612. doi: 10.1016/j.cels.2019.05.005. PubMed PMID: 31202631; PubMed Central PMCID: PMCPMC6857794.

54. Yang Y, Sun W, Liu Y, Zhang J, Li M. An interpretable molecular framework for predicting cancer driver missense mutations. Proc Natl Acad Sci U S A. 2026;123(6):e2524289123. Epub 20260203. doi: 10.1073/pnas.2524289123. PubMed PMID: 41632843; PubMed Central PMCID: PMCPMC12890849.

55. Zhong G, Zhao Y, Zhuang D, Chung WK, Shen Y. PreMode predicts mode-of-action of missense variants by deep graph representation learning of protein sequence and structural context. Nat Commun. 2025;16(1):7189. Epub 20250805. doi: 10.1038/s41467-025-62318-4. PubMed PMID: 40764308; PubMed Central PMCID: PMCPMC12325985.

56. Orenbuch R, Shearer CA, Kollasch AW, Spinner AD, Hopf T, van Niekerk L, et al. Proteome-wide model for human disease genetics. Nat Genet. 2025;57(12):3165–74. Epub 20251124. doi: 10.1038/s41588-025-02400-1. PubMed PMID: 41286104; PubMed Central PMCID: PMCPMC12695638.

